# Ohmline lipid platform: a dual antimicrobial and nanocarrier strategy to potentiate antibiotic efficacy

**DOI:** 10.64898/2026.01.19.699906

**Authors:** Diana Marcos Fernández, Natalia Alfaro, Andrea Cutró, Diego Pazos-Castro, Inés Oliver Camacho, Luis Tebar Palmero, Ana Bouchet, Axel Hollmann

## Abstract

The global rise of antimicrobial resistance has significantly reduced the effectiveness of conventional antibiotics, highlighting the urgent need for alternative and complementary therapeutic strategies. Nanotechnology-based drug delivery systems, particularly lipid nanoparticles, have emerged as promising tools to enhance antibiotic efficacy while limiting toxicity and resistance development. In this study, we evaluated the antimicrobial activity and drug carrier potential of Ohmline, a novel alkyl-ether glycolipid capable of self-assembling into nanotubes and lipid nanoparticles. First, a wide range of Gram-positive and Gram-negative bacteria were used to test Ohmline nanotubes’ antibacterial activity. All examined strains were partially inhibited, with a more noticeable effect on Gram-positive bacteria. Then, the synergistic potential of Ohmline combined with commercially available antibiotics (ampicillin, ceftriaxone, and ciprofloxacin) was evaluated using two different approaches: binary mixtures of Ohmline nanotubes and antibiotics and microfluidically produced Ohmline:DMPC (75:25) nanoparticles with the antibiotics encapsulated. Binary formulations demonstrated strong, strain-dependent synergistic effects at sub-MIC antibiotic concentrations, particularly against *Enterococcus faecalis* and *Citrobacter braakii*. Notably, antibiotic encapsulation within Ohmline nanoparticles further enhanced antimicrobial efficacy compared to non-encapsulated combinations, achieving near-complete growth inhibition in *E. faecalis* and significant inhibition in *Klebsiella pneumoniae* and *C. braakii*.

Overall, our findings demonstrate that Ohmline possesses intrinsic antibacterial activity and acts as an effective lipid nanocarrier that potentiates antibiotic action. The dual functionality of Ohmline supports its potential as a versatile building block for next-generation antimicrobial formulations.

## 1. Introduction

The development of antibiotics marked a major milestone in 20th-century medicine, significantly lowering morbidity and mortality associated with bacterial infections since their use in clinical practice (Baran et al., 2023).

Most antibiotics exert their effects by interfering with essential processes after entering into bacterial cells. However, their extensive and repetitive use in therapy, together with prophylactic use in clinics, have facilitated the emergence of drug-resistant bacteria (Peng et al., 2024).

Antimicrobial resistance is linked to increased mortality rates and elevated healthcare expenses, significantly reducing the effectiveness of antimicrobial treatments. Multidrug resistance (MDR) increases the likelihood of resistant pathogen spread, reduces treatment effectiveness and leads to longer infection durations in patients (Tanwar et al., 2014). The spread of drug-resistant pathogens threatens the ability to treat common infections and perform life-saving procedures, including chemotherapy, organ transplantation, and other surgeries (Alara and Alara, 2024). Moreover, drug-resistant infections have a significant impact on the health of animals and plants and lower agricultural productivity on farms, thus implying a serious risk to global food security (World Health Organization, 2013).

These challenges are further intensified by the inherent difficulties associated with developing new drugs that act through alternative mechanisms. Over the past two decades, only a small number of antibacterial agents have reached approval, and many of them are associated with safety concerns that limit their widespread use. Moreover, the selective pressure imposed by these therapies has often facilitated the rapid emergence and dissemination of resistant bacterial strains shortly after their clinical introduction (González-Freire et al., 2025).

These problematic situations reinforce the necessity of developing new therapeutic alternatives. In this context, nanotechnology-based strategies are being explored to facilitate the concurrent delivery of antibiotics and activation of the immune system. These strategies aim to address the primary challenges associated with antibiotic use: low therapeutic efficacy, long-term toxicity, and the induction of antibiotic resistance (Wani et al., 2024).

It has been demonstrated that inorganic nanoparticles possess bactericidal properties (Singh et al., 2020). Specifically, silver nanoparticles are extensively studied. Despite their bactericidal action, these nanoparticles predominantly accumulate in the lungs and liver and exhibit cytotoxic effects, raising concerns regarding their clinical application. Conversely, lipidic nanoparticles exhibit minimal to no adverse effects and are highly versatile. Therapies using lipidic nanoparticles function as nanocarriers to enhance drug half-life and target specific microorganisms. It has been extensively shown that liposomes improve antibiotic pharmacokinetics and pharmacodynamics and protect the antibiotics against degradation within the body (Eleraky et al., 2020).

Formulations with lipids encapsulating antibiotics have already been approved by the FDA. It’s the case of ARIKAYCE®, which uses dipalmitoyl phosphatidyl choline (DPPC) and cholesterol encapsulating amikacin for *Mycobacterium avium* (Ferreira et al., 2021). Gamal A Shazly (Shazly, 2017) also reported that solid lipid nanoparticles composed of stearic acid had a high percentage entrapment efficiency of ciprofloxacin, and superior antibacterial activity to that of the control. Also, a major drawback commonly reported is the low encapsulation and loading efficiency (Eleraky et al., 2020).

Although these approaches demonstrate an obvious potential, the creation of innovative strategies to combat antibiotic resistance remains essential (Ahmed et al., 2024). Lipids standardized to form nanoparticles (such as DOPC, DPPC, DPTAP, DOPE, DMPG, cholesterol) lack intrinsic bactericidal activity. In most studies, the empty lipid nanoparticle was used as a negative control of activity and induced no bacteria depletion.

1-O-hexadecyl-2-O-methyl-sn-glycero-3-β-lactose (Ohmline) is an alkyl-ether lipid with a lactose group as the polar head and a hydrophobic tail (Girault et al., 2011). It was previously described that Ohmline has the ability to partition into the lipid membrane and modify its physicochemical properties (Herrera et al., 2017). These investigations suggest that Ohmline could act through local modulation of the biophysical properties of the plasma membrane (Herrera et al., 2017), and may inhibit bacteria by altering the properties of its membranes. In addition, it was previously demonstrated that Ohmline has the capability to spontaneously form nanotubes or lipidic nanoparticles (Villanueva et al., 2024). This implies that, in addition to its potential antimicrobial activity, Ohmline could also function as an antimicrobial carrier.

In this article, we aimed to evaluate the potential use of Ohmline as an antimicrobial agent and a lipid nanocarrier. A wide spectrum of bacterial strains was analyzed to determine the effect of Ohmline, including Gram-positive and Gram-negative organisms. The Gram-positive organisms analyzed were *Enterococcus faecalis*, *Bacillus subtilis*, *Lactococcus* spp., and *Staphylococcus aureus*. The Gram-negative organisms analyzed were *Pseudomonas aeruginosa*, *Klebsiella pneumoniae*, and *Escherichia coli*. In addition, the synergy of binary formulations of Ohmline and different antibiotics was evaluated. Finally, given its potential as a lipidic nanocarrier, Ohmline was also used to encapsulate several antibiotics to evaluate the inhibition of bacterial growth with this combined therapy.

## 2. Material and methods

### 2.1 Materials

DMPC (1,2-dimyristoyl-sn-glycero-3-phosphocholine) was obtained from Avanti Po-lar Lipids (USA). Ohmline was obtained from Lifesome Therapeutics (Spain). The water used was ultra-pure (conductivity: 0.002 ± 0.001 μS/cm; pH= 5.0). The organic solvents used (chloroform, methanol) were all analytical grade, purchased from Merck Química SAIC (Argentina). For the microbiological tests, Mueller Hinton broth (MHB) was obtained from Oxoid (England). The antimicrobial agents used, ampicillin sodium salt (AMP) and ceftriaxone sodium salt hemiheptahydrate (CEF) were purchase on Thermo Fisher Scientific (USA) and ciprofloxacin (CFX) was purchase on MedChemExpress (USA).

### 2.2 Bacteria Strains

The reference strains *Bacillus subtilis* ATCC 6633, *Escherichia coli* ATCC 25922, *Pseudomonas aeruginosa* ATCC 27853, and *Staphylococcus aureus* ATCC 2593 were obtained from commercial suppliers. *Enterococcus faecalis*, *Lactococcus* spp., and *Citrobacter braakii*, were isolated from animal-derived samples collected at a slaughterhouse in Spain. Bacterial colonies were isolated and identified by MALDI-TOF. These strains were generously supplied by the research group led by Christian Ghidelli at Funditec. The *Klebsiella pneumoniae* strains Kpn10 and Kpn11 were a clinical isolates obtained from patients admitted to Hospital Universitario Ramón y Cajal, Madrid, Spain (Calvo-Villamañán et al., 2025) and was kindly provided by the Centro Nacional de Biotecnología (CNB-CSIC), Madrid, Spain.

### 2.3 Ohmline Stock Solutions

Stock solutions of Ohmline (Lifesome Therapeutics) were prepared at concentrations of 5 mg/ml and 10 mg/ml. The compound was weighed in glass vials using a precision balance and dissolved in sterile high-performance liquid chromatography (HPLC)-grade water.

### 2.4 Preparation of Antibiotics Encapsulated by Microfluidic method

Ohmline and DMPC were weighed separately in glass vials using a precision balance. DMPC was dissolved in chloroform to a final concentration of 5 mg/ml. Ohmline was dissolved in a chloroform:methanol mixture (8:2 v/v) to the same final concentration (5 mg/ml), assisted by 10 minutes of sonication. Using a Hamilton syringe, the two solutions were combined to achieve the desired 75:25 molar ratio in the organic phase. The solvent was then evaporated with nitrogen stream, forming a lipid film that was subsequently resuspended in high precision liquid chromatography (HPLC)-grade water.

The resulting lipid suspension underwent five cycles of sonication followed by heating (10 minutes each). Nanoparticles containing Ohmline:DMPC (75:25) encapsulating antibiotics were produced using the TAMARA microfluidic formulation system (Inside Therapeutics, France) under sterile conditions with a Bunsen burner flame. All runs were performed using a herringbone microfluidic chip, with a total flow rate (TFR) of 12,000 µl/min and a flow rate ratio (FRR) of 10:1 (aqueous phase:lipid solution). From each run, 1 ml of nanoparticle suspension was obtained in sterile Eppendorf tubes. Between runs, the system was cleaned with water and/or ethanol.

### 2.5 Nanoparticle characterization

Ohmline:DMPC (75:25) at a concentration of 0.5 mg/ml with different antibiotic concentrations obtained with the microfluidics in HPLC-grade water were prepared. Samples were then diluted to 0.1 mg/ml prior to analysis. Particle size and zeta potential were measured using dynamic light scattering (DLS) with a ZetaSizer Nano ZS analyzer (Malvern Instruments Ltd, UK)

### 2.6 Minimum inhibitory concentration

Minimum inhibitory concentrations (MICs) were determined for each antibiotic and bacterial strain using the broth microdilution method, following the guidelines established by the Clinical and Laboratory Standards Institute (CLSI) (Limbago, 2024). Briefly, an aliquot of an overnight culture was diluted in MHB to obtain an inoculum of approximately 1 × 10⁶ CFU/ml. Stock solutions of the antibiotics were prepared under sterile conditions and subsequently subjected to serial two-fold dilutions in 96-well microtiter plates, covering the concentration ranges recommended by CLSI for each bacterial group evaluated. The microplates were incubated at 37 °C for 18–24 h under aerobic, static conditions. The MIC was defined as the lowest antibiotic concentration that inhibited visible bacterial growth. To ensure reproducibility, all assays were performed in triplicate in at least three independent experiments.

### 2.7 Plaque Assay

In 96-well polystyrene plates (U-bottom), different experimental conditions were evaluated: a) *Batch assay:* Ohmline at 125 µg/ml, 250 µg/ml, and 500 µg/ml was evaluated as monotherapy, or sequentially combined with different concentrations of AMP, CEF, or CFX b) *Microfluidics assay,* where Ohmline:DMPC (75:25) at 250 µg/ml and 500 µg/ml was combined with the same antibiotics but at different concentrations. All assays were compared with their respective negative controls. After preparation, the bacterial inoculum incubated in MHB was added to all wells except for negative controls of MHB only. Growth controls were also included containing bacteria and MHB. The effect of the combinations on the growth curves of *E. faecalis*, *C. braakii*, *E. coli, S. aureus* and *K. pneumoniae* KPN10 was then evaluated under the corresponding conditions for each well. The final volume in each well was 220 µl, and the bacterial concentration was standardized to approximately 10⁴ CFU per well. Absorbance at 600 nm was measured every 30 minutes over a 48-hour period at 37°C, with continuous orbital shaking throughout the assay in a Microplate reader (SPECTROstar Omega, BMG LABTECH, Germany).

### 2.8 Inhibition of biofilm formation

The inhibition of biofilm development was assayed in 96-well flat-bottom polystyrene plates. Briefly, the Ohmline was serially diluted in MHB, after which the cell suspensions were inoculated into the wells to a final concentration of 5 × 10^5^ CFU/ml. The plates were incubated at 37 °C for 72 hrs. In addition, MHB was added to blank wells without bacterial culture. After the incubation period, the wells were washed three times with sterile 20 mM PBS (pH= 7.40) to remove nonadherent cells. The microplates were dried at 60 °C for 1 h to fix the biofilm. The wells were stained with 0.5% (w/v) crystal violet and incubated at room temperature for 15 min. Later, the plates were washed six times with sterile distilled water to remove unabsorbed stains and dried at 37 °C for 30 min. Each well was dissolved in ethanol at 96%, and the absorbance was measured at OD_595_ nm using a microplate reader (ELx808iu, BioTek, USA). The percentage inhibition was calculated following Equation (Bazargani and Rohloff, 2016; Merghni et al., 2016).

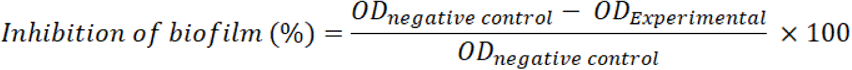

### 2.9 Statistical analysis

Most experiments were performed with three or four independent measurements, and results are presented as means with error bars representing either standard deviation (SD) or standard error of the mean (SEM) of the averaged values. Data were analyzed using R and GraphPad Prism software. Statistical significance was determined using one-way ANOVA followed by Tukey’s multiple comparisons test.

## 3. Results

### 3.1 Antibacterial effect of Ohmline nanotubes

As a first step, the antibacterial activity of Ohmline nanotubes was tested by following the optical density of the culture. Fig. 1 shows that Ohmline can, at least partially, inhibit bacterial growth of all strains tested. The best results were observed with the Gram-positive strains, with inhibition scores ranging from 28 to 45%. In the case of Gram-negative strains, apart from *E. coli* the inhibition found was below 20%, suggesting a lower effect on Gram-negative bacteria. A control growth curve of *S. aureus* with DMPC at 500 µg/ml was carried out without and it did not diminish the bacterial growth (Fig. S1).

**Figure 1:**
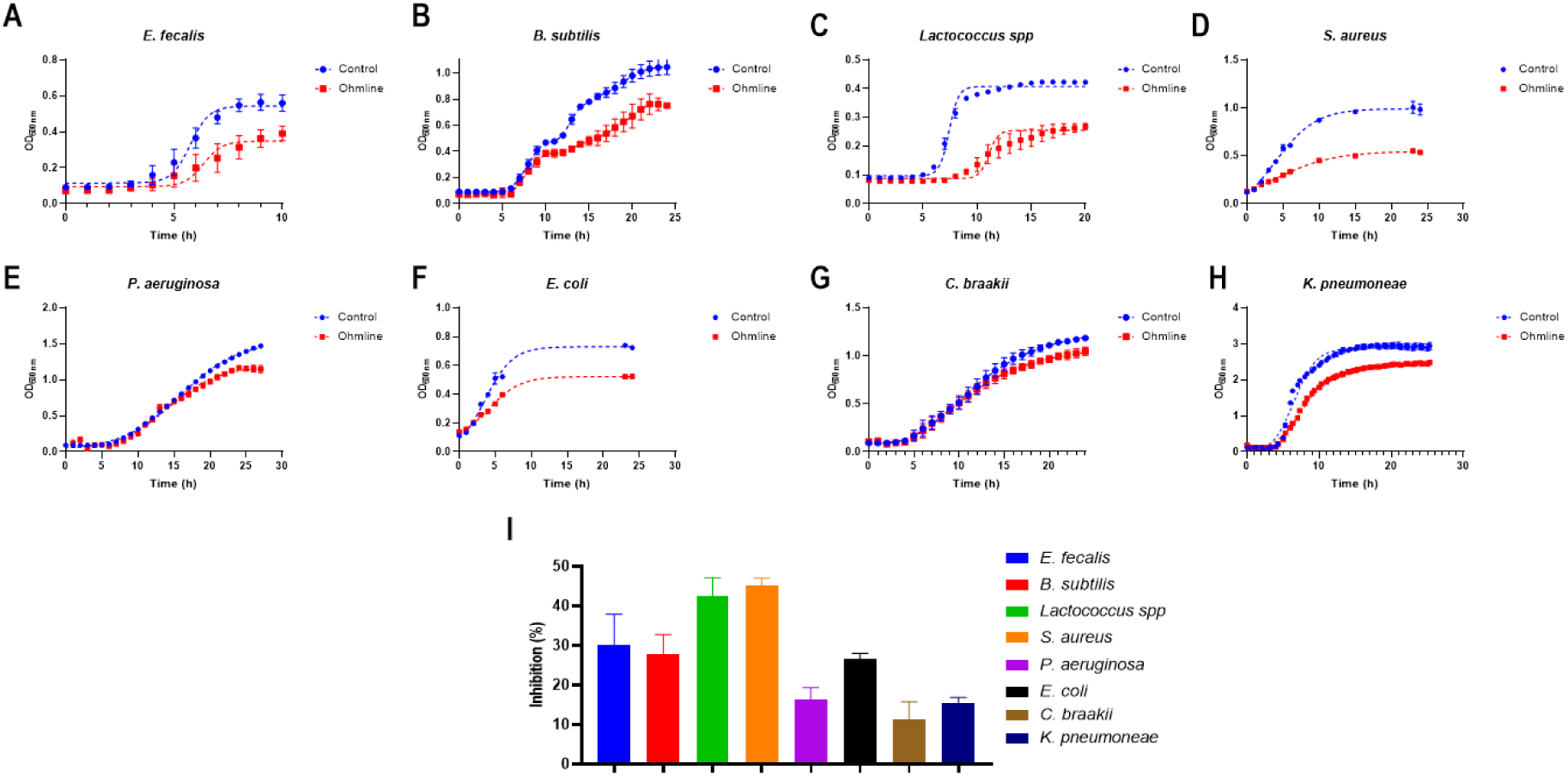
Antibacterial effect of 500 ug/ml of Ohmline nanotubes. Growth curves for **(A)** Enterococcus faecalis, **(B)** Bacillus subtilis, **(C)** Lactococcus spp, **(D)** Staphylococcus aureus, **(E)** Pseudomonas aeruginosa, **(F)** Escherichia coli, **(G)** Citrobacter braakii and **(H)** Klebsiella pneumoniae in the absence and presence of Ohmline 500 μg/ml. Percentage of inhibition induced for Ohmline at final time for all bacteria strains. **(I)**. The data represents an average of three independent measurements; the error bars indicate the standard deviation of the averaged values.

Following the promising results on *S. aureus*, and as proof of concept, its effect on inhibition of 72-h biofilm formation was also evaluated. A reduction in biofilm mass of approximately 40% was observed (Fig. S2).

### 3.2 Effect of Ohmline in combination with commercially available antibiotics

The potential of Ohmline as a carrier or booster when combined with commercially available antibiotics was then evaluated. As a first step, binary formulations of Ohmline nanotubes and several concentrations of commercial antibiotics AMP and CEF were analyzed.

#### 3.2.1 Binary combinations of Ohmline nanotubes with antibiotics

Before conducting the assays with the binary formulations, the MIC values for each antibiotic was determined for each bacteria obtaining MIC values in line with previously described in literature (Table S1).

As shown in Fig. 2, the addition of Ohmline at sub-MIC concentrations of the antibiotics significantly enhances the inhibitory effect in *E. faecalis* and *C. braaki.* In the case of *S. aureus,* the effect is observed at sub-MIC concentrations close to the MIC. The highest scores of synergism for each bacterium tested depend on both Ohmline and antibiotic concentration (Table S2). Both AMP and CEF showed a high zero interaction potency (ZIP) synergy scores (9.48 and 16.61 respectively) with Ohmline for the *E. faecalis* strain (Fig. S3), indicating synergic interaction.

**Figure 2.**
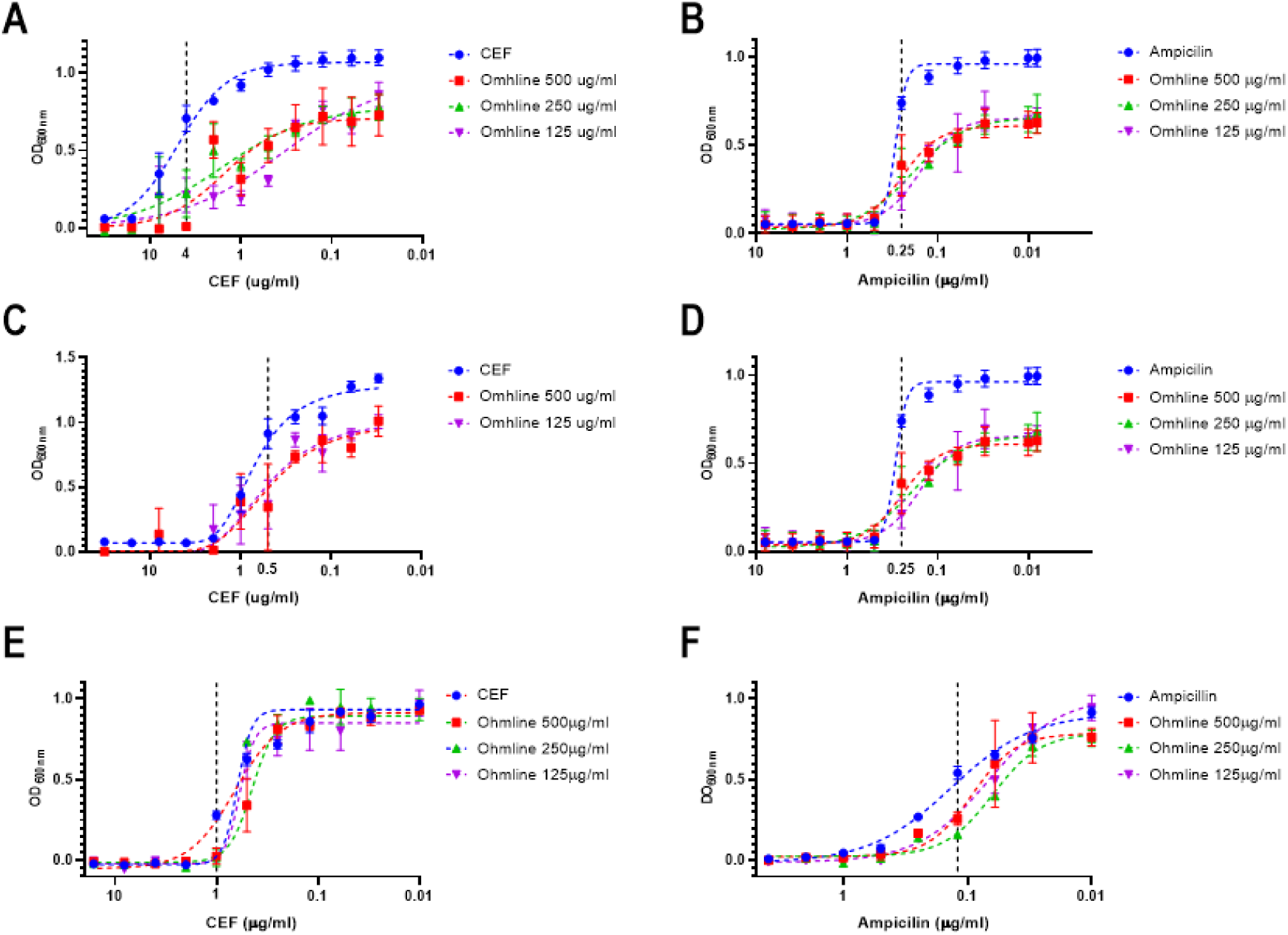
Effect of binary combination of Ohmline and ceftriaxone (CEF) or ampicillin. The effect of binary combination is tested in *Enterococcus faecalis* **(A, B),** *Citrobacter braakii* **(D, C)** and *Staphylococcus aureus* **(E, F)** Optical density was represented as a function of antibiotic concentration and a non-linear regression fit (variable slope) was calculated. The data represent an average of four independent measurements, with four technical replicates in the case of the antibiotic curve; the error bars indicate the standard deviation of the averaged values.

To highlight the results obtained, data of Fig. 2 were plotted as inhibition percentage data showing the results for one binary formulation for each bacterium and antibiotic tested (Fig. S4).

It can be concluded that the addition of Ohmline to sub-MIC concentrations of the antibiotics synergistically increases the antimicrobial activity, resulting in at least 75% inhibition of bacterial growth after 24 hours of incubation. Furthermore, in the case of *S. aureus,* almost complete inhibition was reached at half the MIC of CEF with 125 µg/ml of Ohmline (Fig. S4).

Interestingly, no significant effects on bacterial growth were recorded in *E. coli* after 24 h of incubation when AMP was combined with Ohmline at 500 or 125 µg/ml (Fig S5).

#### 3.2.2 Antibiotics encapsulated on Ohmline formulations

To evaluate the feasibility of using Ohmline as an antibiotic carrier, Ohmline-based nanoparticles encapsulating antibiotics were obtained using a microfluidic method (Pittiu et al., 2024). DMPC at 25% (w/w) was included in the formulation to enhance the stability, as it is well known that DMPC is stable to oxidation and readily hydrates in water, forming lamellar phases at physiological pH and temperature (Shireen et al., 2015).

First, DLS characterization of empty Ohmline:DMPC (75:25) nanoparticles was assessed and showed that most of the particles obtained (> 80 %) exhibited a size distribution between 60 - 200 nm (Fig. S6). However, a small proportion of particles with larger sizes were recorded, resulting in a polydispersity index (PDI) of approximately 0.8. Regarding surface charge, as expected, the zeta potential of the empty formulations exhibited values of +0,443 mV (Fig. S7-A). Encapsulation of CFX enhanced the nanoparticle stability, with a recorded Z-average of 205.4 ± 3.3 nm and PDI 0.39 for 0.25 µg/ml. Similarly, the addition of CEF also stabilize the lipid nanoparticles, measuring a Z-average of 155.0 nm and PDI 0.37 when combined with Ohmline:DMPC. In terms of surface charge, nanoparticles loaded with CFX showed a mean negative Z potential (−3.5 mV); while CEF-loaded nanoparticles registered a mean positive Z potential (+7.3 mV). The addition of AMP to the Ohnline:DMPC nanoparticles did not alter nanoparticle stability nor surface charge (Fig. S6).

Then, we investigated the effect those antibiotics-loaded Ohmline:DMPC nanoparticles on bacterial growth. For almost all strains and conditions tested, an enhanced inhibitory effect was observed compared to the same dose of free antibiotic. (Table S3).

In *E. faecalis,* encapsulation of different sub-MIC concentrations of AMP with Ohmline:DMPC enhances the effect of the antibiotic tested, either by increasing the lag phase or resulting in a lower bacterial mass at the end of incubation (Fig. 3 A-B; Fig S.8 A-B). Remarkably, at the highest AMP concentration tested (half of the MIC), a highly synergistic effect was observed with 500 µg/ml of nanoparticles (i.e. 372 µg/ml of Ohmline), as no bacteria growth was detected after more than 24 h of incubation.

**Figure 3.**
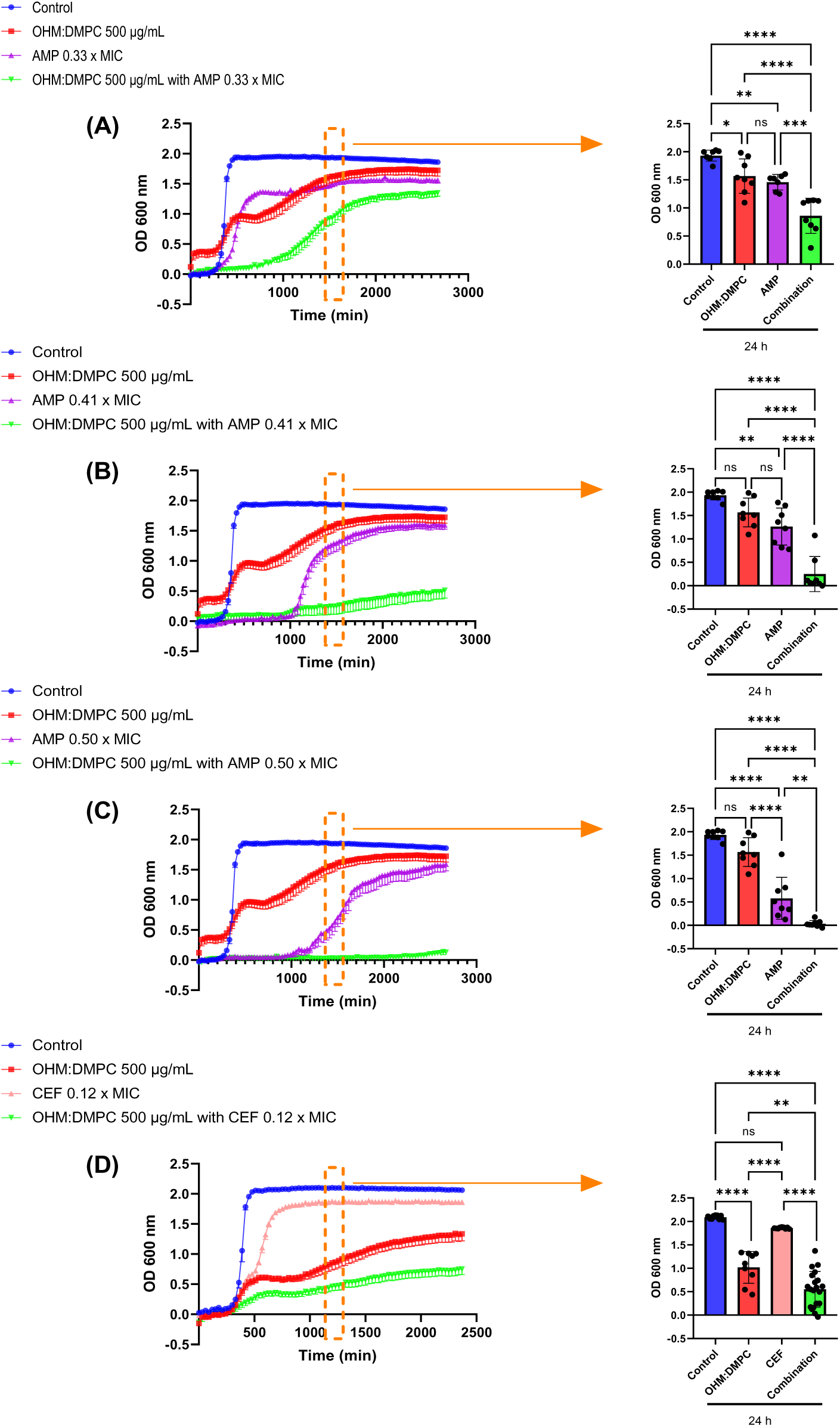
Effect of encapsulation of antibiotics on Ohmline nanoparticles at 500 µg/m concentration on the growth of *Enterococcus faecalis*. Growth curves of *Enterococcus faecalis* treated with ampicillin (AMP) **(A–C)** or ceftriaxone (CEF) **(D)** encapsulated in Ohmline:DMPC (75:25) at a concentration of 500 μg/ml. Data are presented as the mean of three independent experiments with 3–4 technical replicates for ampicillin and 3–7 technical replicates for ceftriaxone. Error bars indicate - SEM. Statistical analysis was performed using one-way ANOVA followed by Tukey’s multiple comparisons test. Statistical significance is indicated as p < 0.05 (*), p < 0.01 (**), p < 0.001 (***), and p < 0.0001 (****). For each antibiotic, the same control growth curve was used for all concentrations.

When the encapsulation of CEF at 0.12 x MIC on Ohmline:DMPC (75:25) was evaluated on *E. faecalis*, a significant increase in growth inhibition was observed (Fig. 3 D; Fig. S8 D). For the highest concentration of Ohmline:DMPC tested, an inhibition of around 65% was reached, again showing a stronger effect than the binary combination of Ohmline and CEF at the same concentration.

An additive effect of CEF encapsulated in Ohmline:DMPC (75:25) at both concentrations tested (0.01 and 0.015 x MIC) was observed in *C. braakii*, at the highest concentration of lipid nanoparticles measured (500 µg/ml) (Fig 4B - E). In the same way, CEF at 0.015 x MIC on 250 µg/ml of nanostructures also showed a significant increase in the antimicrobial activity (Fig. 4D). Whereas the lowest concentration of CEF (0.01 x MIC) encapsulated in 250 µg/ml of liposomes has a statistical significance, but lower inhibition effect (Fig. 4A).

**Figure 4.**
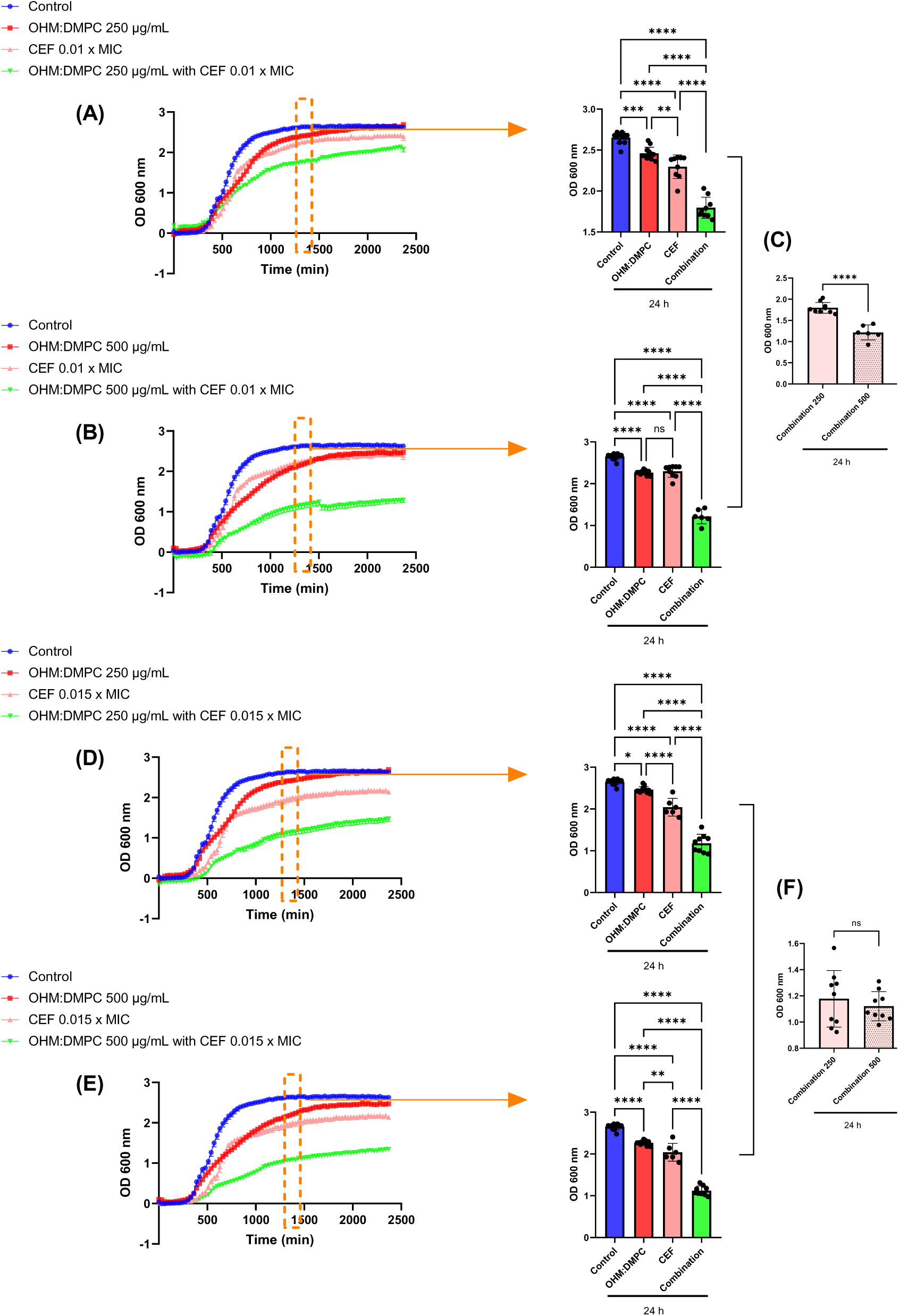
Effect of encapsulation of ceftriaxone on Ohmline nanoparticles on the growth of *Citrobacter braakii*. Growth curves of *Citrobacter braakii* treated with ciprofloxacin (CFX) at different concentrations combined with Ohmline:DMPC at 250 μg/ml **(A, D)** or 500 μg/ml **(B, E).** Comparison of bacterial growth at 24 h following treatment with CFX at 0.01× MIC combined with Ohmline:DMPC at 250 or 500 μg/ml **(C)**. Comparison of bacterial growth at 24 h following treatment with CFX at 0.015× MIC combined with Ohmline:DMPC at 250 or 500 μg/ml **(F)**. Data are presented as the mean of four independent experiments with 3–6 technical replicates each. Error bars indicate - SEM. Statistical analysis was performed using one-way ANOVA followed by Tukey’s multiple comparisons test. Statistical significance is indicated as p < 0.05 (*), p < 0.01 (**), p < 0.001 (***), and p < 0.0001 (****). The same control growth curve was used for all treatment combinations.

For *S. aureus*, two antibiotics were tested: CEF and CFX. For the first antibiotic, a significant effect was observed at both concentrations of Ohmline:DMPC (75:25) after 24 hours of incubation, however, this effect diminished over the incubation period (Fig. 5A-B). For the second antibiotic (CFX) both concentrations of Ohmline:DMPC did also produce a significant reduction in the bacterial growth rate (Fig. 5 D-E). In this case, the major effect was observed as an extended lag phase, indicating a stronger inhibition of early bacterial proliferation.

**Figure 5.**
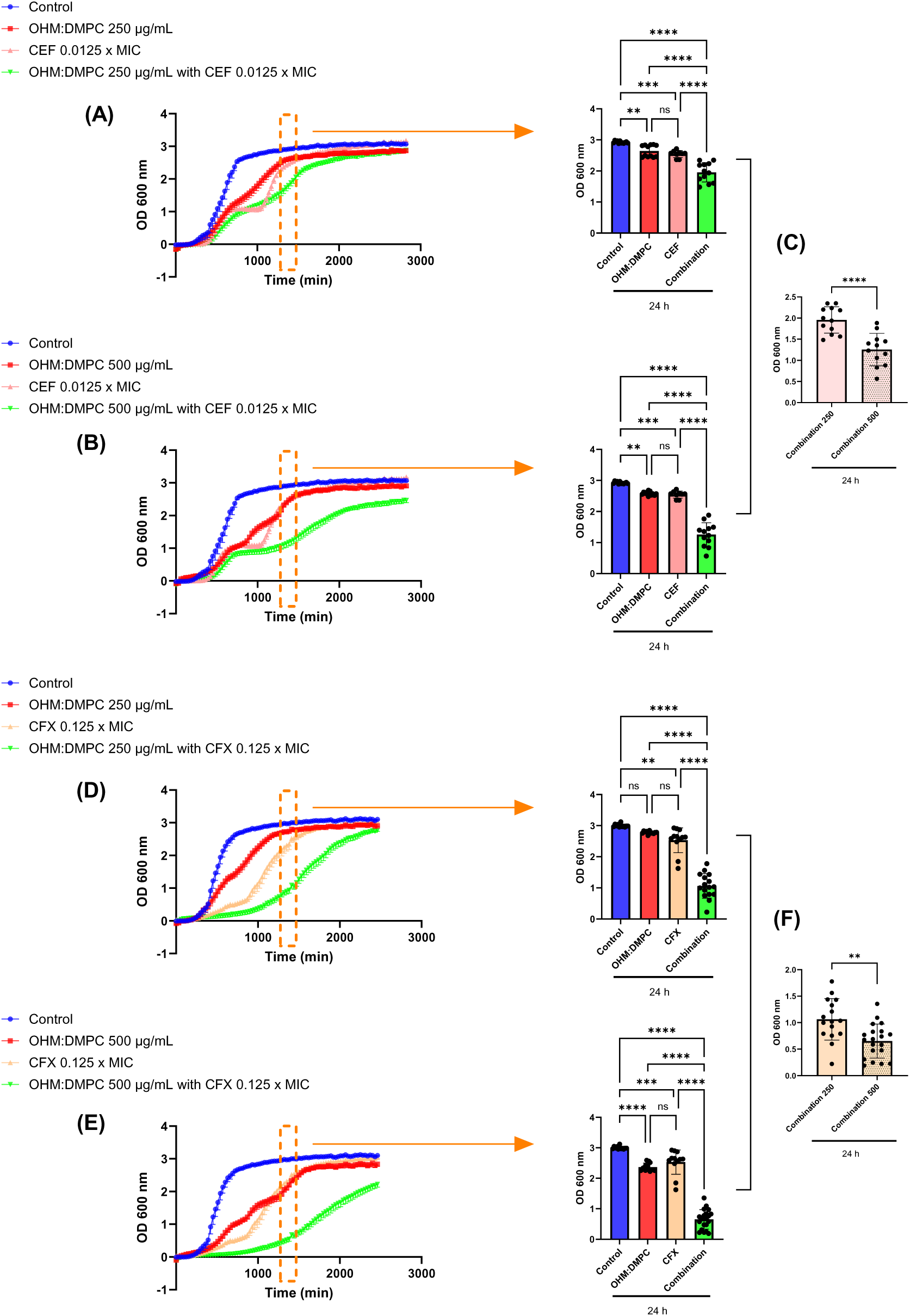
Effect of antibiotic encapsulation in Ohmline nanoparticles on the growth of *Staphylococcus aureus*. Growth curves of *Staphylococcus aureus* treated with ceftriaxone (CEF) at 0.0125 × MIC combined with Ohmline:DMPC at 250 μg/ml **(A)** or 500 μg/ml **(B)**. Comparison of bacterial growth at 24 h following treatment with CEF combined with Ohmline:DMPC at 250 or 500 μg/ml **(C).** Growth curves of *Staphylococcus aureus* treated with ciprofloxacin (CFX) at 0.125 × MIC combined with Ohmline:DMPC at 250 μg/ml **(D)** or 500 μg/ml **(E)**. Comparison of bacterial growth at 24 h following treatment with CEF combined with Ohmline:DMPC at 250 or 500 μg/ml **(F).** Data are presented as the mean of 3–4 independent experiments with three technical replicates each. Error bars indicate - SEM. Statistical analysis was performed using one-way ANOVA followed by Tukey’s multiple comparisons test. Statistical significance is indicated as p < 0.01 (), p < 0.001 (***), and p < 0.0001 (****). The same control growth curve was used for all graphs corresponding to the same antibiotic.

In the case of *E. coli*, in the same line with previous assays, no synergistic effects were observed using different AMP concentrations encapsulated in 250 or 500 µg/ml of Ohmline:DMPC nanoparticles (Fig. S7)

Finally, the effect of the encapsulation of CFX at 0.12 x MIC in Ohmline:DMPC (75:25) nanoparticles was evaluated on *K. pneumoniae* (Fig. 6). In both strains at 500 µg/ml of Ohmline:DMPC, almost 50% of inhibition was recorded after 40 h of incubation. This inhibition was significantly higher than the 15% obtained for Ohmline:DMPC (75:25) or CFX alone. On the other hand, at 250 µg/ml of Ohmline:DMPC, only a slight difference was obtained in comparison with CFX or Ohmline:DMPC (75:25) alone.

**Figure 6.**
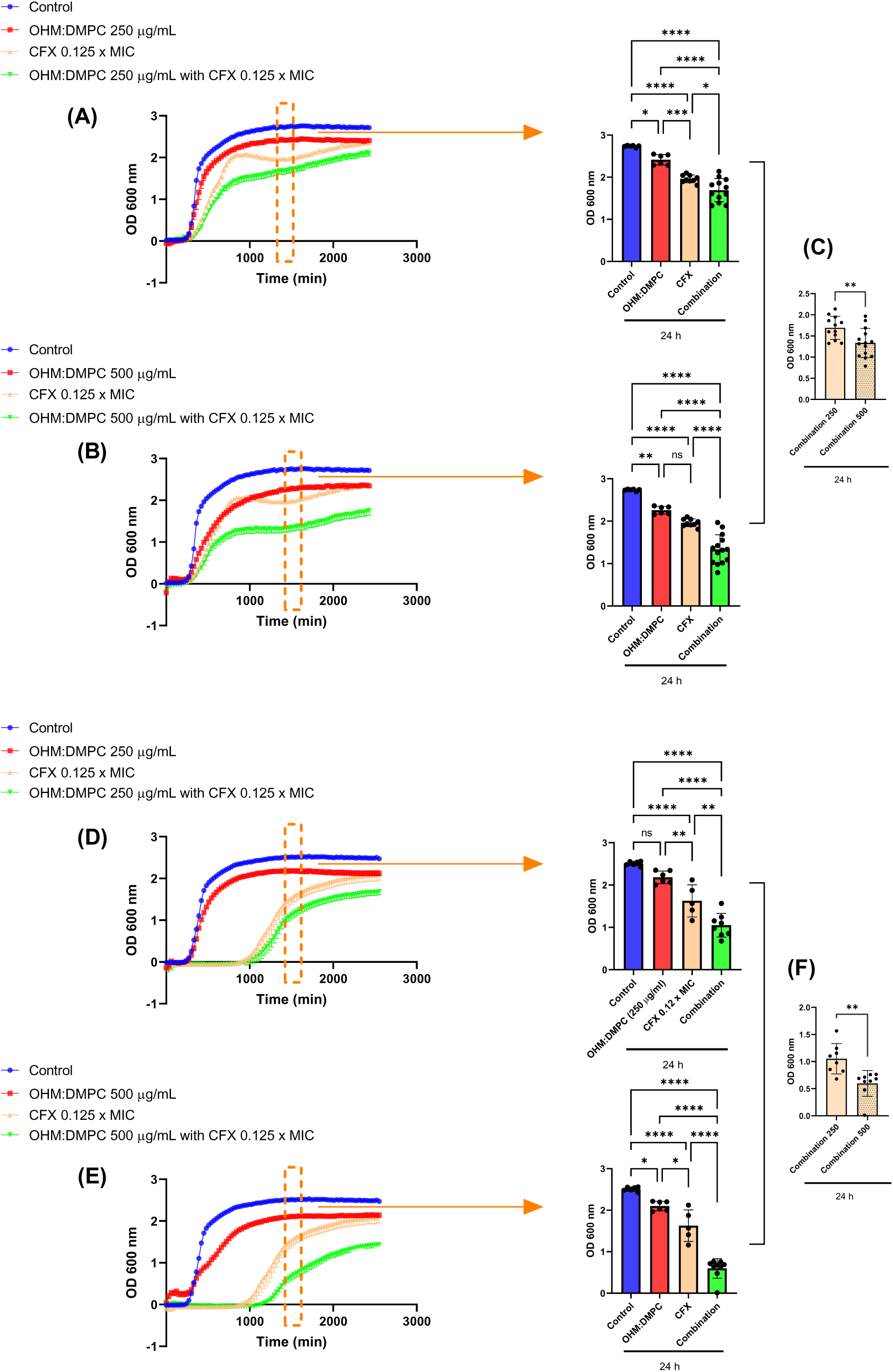
Effect of encapsulation of ciprofloxacin on Ohmline nanoparticles on the growth of *Klebsiella pneumoniae*. Growth curves of *Klebsiella pneumoniae* Kpn10 treated with ciprofloxacin (CFX) combined with Ohmline:DMPC at 250 μg/ml **(A)** or 500 μg/ml **(B)**. Comparison of bacterial growth at 24 h for Kpn10 treated with CFX combined with Ohmline:DMPC at 250 or 500 μg/ml **(C).** Growth curves of a clinical *K. pneumoniae* isolate treated with CFX combined with Ohmline:DMPC at 250 μg/ml **(D)** or 500 μg/ml **(E).** Comparison of bacterial growth at 24 h for the clinical isolate treated with CFX combined with Ohmline:DMPC at 250 or 500 μg/ml **(F)**. Optical density over time is shown as mean - SEM. Bar graphs represent mean optical density values at 24 h. Two independent experiments with 3–9 technical replicates each were performed. Statistical analysis was conducted using one-way ANOVA followed by Tukey’s multiple comparisons test. Statistical significance is indicated as p < 0.05 (*), p < 0.01 (**), p < 0.001 (***), and p < 0.0001 (****). For each bacterial strain, the same control curve was used for comparisons with Ohmline:DMPC at both concentrations.

## 4. Discussion

There is an urgent need to develop innovative antimicrobial therapies against persistent bacterial infections. Although the main focus of efforts has been on developing new chemical agents, in the current pressing scenario, repurposing and reformulating already-existing molecules is thought to be a speedier strategic strategy. As a result, current efforts to develop novel antimicrobial treatments have focused on enhancing the way that commercially available antibiotics are delivered. In particular, lipids show great promise as building blocks for drug delivery system design. Possessing innate antibacterial qualities and the capacity to self-assemble into structural drug delivery vehicles (Thorn et al., 2021).

In the present work, the antimicrobial and potential carrier abilities of a novel alkyl-ether lipid called Ohmline were studied. Ohmline nanotubes showed on several bacteria an inhibitory activity but with a more pronounced effect on Gram-positive ones. Previously, it was described that Ohmline has the ability to partition into the lipid membranes and modify its physicochemical properties (Herrera et al., 2017). Therefore, it was postulated that these perturbations induced in the bacterial membrane could compromise its viability.

The differential trend on Gram-negative and Gram-positive bacteria could be explained by the differences in their bacterial envelopes. Gram-negative bacteria have two membranes that make up their cell envelope: the outer membrane (OM) is asymmetric and primarily formed of lipopolysaccharides, while the inner, cytoplasmic membrane (IM) is made up of a phospholipid bilayer. A layer of peptidoglycan is found in the periplasmic region, which lies between the two membranes (Schwechheimer and Kuehn, 2015). Although Gram-positive bacteria do not have an OM, they are encased in peptidoglycan layers that are many times thicker than those of Gram-negative bacteria (Savini et al., 2020). It was previously reported that Carbohydrate fatty acid derivatives have been shown to have better inhibitory effects on Gram-positive rather than Gram-negative bacteria (Piao et al., 2006; Wagh et al., 2012). In the same line, it was pointed out that Gram-negative bacteria were more resistant to Carbohydrate fatty acid derivatives due to lipopolysaccharides present in their outer membrane (Zhao et al., 2015).

Ohmline has the unique characteristic of spontaneously forming nanostructures (Villanueva et al., 2024). This characteristic allowed us to consider strategies where Ohmline could also act as a drug carrier, which could be further enhanced by its inherent, albeit moderate, antimicrobial activity. In this context, the potential of Ohmline when combined with commercially available antibiotics was evaluated. For a better comprehension of the feasibility of Ohmline as a carrier, as a first step, binary formulations of Ohmline and the commercial antibiotics AMP and CEF were evaluated, to later inquire about formulations in which the antibiotics were encapsulated in Ohmline nanoparticles.

For most of the combined formulation, a synergistic or additive effect was found at sub-MIC antibiotic concentrations; this effect showed strong dependence on the antibiotics used and strain tested. In line with that, other studies that evaluate synergic formulations with antibiotics also reported a highly strain-dependent behaviour (Buyck et al., 2015; Valderrama et al., 2020). These results strongly support the potential of Ohmline to enhance antimicrobial action of other agents.

To particularly evaluate the feasibility of using Ohmline as an antibiotic carrier, a second combined formulation of Ohmline with AMP or CEF was evaluated, but in this case, the antibiotics were encapsulated into Ohmline nanostructures, which also contained DMPC to provide greater structural stability, using the microfluidic method. It should be pointed out that pure DMPC liposomes did not show any antibacterial effect, highlighting the specific contribution of Ohmline to bacterial inhibition.To investigate another group of antibiotics, CFX was also tested in this step in *K. pneumoniae* and *S. aureus*.

The physicochemical characterization of Ohmline:DMPC with DLS demonstrated that empty formulations contained nanoparticles with a diameter between 60 - 200 nm and a small number of large particles or nanotubes. It should be marked that Z potential showed almost neutral surface charge. It was previously described that Ohmline is a neutral lipid (Villanueva et al., 2024), and DMPC is well known as a zwitterionic lipid with no net charge (Shireen et al., 2015). The incorporation of antibiotics could influence the physicochemical properties of the nanostructures. In this way, the encapsulation of CFX or CEF resulted in improved nanoparticle stability. On the other hand, AMP did not induce a comparable stabilization effect. The Z potential was also demonstrated to be dependent on the antibiotic used to formulate the nanoparticle. These observations highlight the sensitivity of the Ohmline nanostructure to drug-lipid interactions and suggest that tuning the lipid composition or Ohmline:lipid ratios could further optimize nanoparticle performance.

These findings are consistent with key design principles of lipid-based drug delivery: the therapeutic behaviour of liposomes and DMPC-derived nanoparticles is governed by lipid identity, membrane rigidity, surface charge, manufacturing method, and particle size. Nanoparticles smaller than 200 nm are generally preferred in biomedical applications because they reduce immune clearance and facilitate penetration through biological barriers. Low PDI and high stability are essential to ensure reproducible release and robust therapeutic efficacy (Jaradat et al., 2022). In this light, the ability of Ohmline to generate stable, antibiotic-loaded nanostructures underscores its promise as a versatile lipid scaffold for advanced antimicrobial formulations. Other formulations are currently being developed to increase the stability and PDI of the nanostructures.

Regarding the antimicrobial properties of antibiotics loaded in Ohmline:DMPC formulations, as for binary combinations, the effects depend on both the antibiotic encapulated and the strain tested. The most remarkable effects were found for *E. faecalis,* where full inhibition for more than 24 h was observed when half of the MIC of AMP was encapsulated on high concentration of Ohmline:DMPC. It should be pointed out that this effect was greater than the binary combination of Ohmline at 500 µg/ml with AMP, indicating that this method significantly enhances antimicrobial action, reinforcing the potential of Ohmline as a drug carrier. Remarkable inhibitions were also found for *K. pneumoniae* and *C. braakii*.

In addition to improving stability, bacterial uptake and solubility, antibiotic encapsulation can direct the drug selectively to infection sites, increasing local concentration and limiting exposure of healthy tissues (Gonzalez Gomez and Hosseinidoust, 2020). When the delivery system exhibits antibacterial activity *per se,* besides the previous points, a synergistic effect by the combination of two antimicrobials could also take place.

The setup of the experiments carried out in this work, where formulations were used as soon as they were obtained, allows us to speculate that an improvement in stability should not be the main point of the increased antimicrobial action observed. On the other hand, the potential interaction of Ohmline with the bacterial envelope could enhance the delivery of antibiotics into the bacteria. It is well established the concentrating effects of membrane-associated drugs, lateral diffusion of drugs across two-dimensional surface (rather than three dimensions in aqueous bulk) could increase reaction rates with receptors in a mechanism referred to as “reduction of dimensionality” (McCloskey and Poo, 1986; Sykes et al., 2014). Furthermore, the ability of Ohmline to alter the membrane properties could also favor the internalization of the antibiotics, explaining the enhanced activity observed. In this regard, it was previously postulated that the self-assembled glycolipids could span through the structurally alike bacterial cell membrane and thereby facilitate the entry of antibiotics (Joshi-Navare and Prabhune, 2013).

## 5. Conclusion

Overall, the antimicrobial activity *per se* of Ohmline, in addition to the enhanced antimicrobial action of commercial antibiotics in both, binary formulation and encapsulated on Ohmline nanoparticles, allows us to postulate Ohmline as a novel molecular scaffold to be considered for antimicrobial applications. Furthermore, the ability of Ohmline to adopt different nanostructures as liposomes or nanotubes, or hybrid nanoparticles reinforces the potential of Ohmline as a novel building block for novel antimicrobial formulations.

## 6. Acknowledgements

The authors acknowledge the financial support of CONICET (PIP 112202101 00756CO), Universidad Nacional de Santiago del Estero (PI-UNSE 23/E012-Bint-2023, PI-UNSE 23/A300), Ministerio de Ciencia, Innovación y Universidades (Spain) and by the Centro para el Desarrollo Tecnológico y la Innovación (CDTI), within the framework of the call “MULTI-PAIS” 2024 (PAIS-20241089). AC, AH are members of the Research Career of CONICET.

