## Supplementary data for "Ohmline lipid platform: a dual antimicrobial and nanocarrier strategy to potentiate antibiotic efficacy"

**An Ohmline-based nanoplatform to reformulate and potentiate antibiotics efficacy**

**^Supplementary Material^**

**Supplementary Tables**

**Supplementary Table 1: Minimum inhibitory concentration of bacteria at 24 h.** The minimum inhibitory concentration for each bacteria was obtained through plaque dilution method. The maximum dose tested was 64 or 128 μg/ml

|  | Ampicillin (µg/ml) | Ceftriaxone (µg/ml) | Ciprofloxacin (µg/ml) |
| --- | --- | --- | --- |
| *Enterococcus faecalis* | 6 | 8 | Not tested |
| *Citrobacter braakii* | Not tested | 11,5 | Not tested |
| *Staphylococcus aureus* ATCC 25922 | 0.5 | 2 | Not tested |
| *Staphylococcus aureus* USA300 | 60 | 64 | 32 |
| *Escherichia coli* | 7 | Not tested | Not tested |
| *Klebsiella pneumoniae* Kpn10 | Not tested | Not tested | 2 |
| *Klebsiella pneumoniae* clinical sample | Not tested | Not tested | 0.5 |

**Supplementary Table 2:** Synergy scores of Ohmline combined with Ampicillin or Ceftriaxone in *Citrobacter braakii, Enterococcus faecalis* and *Staphylococcus aureus*. Obtained with Synergyfinder software (<https://synergyfinder.aittokallio.group/2025110612260025589/>), last visited 31/07/2025.

| Drug combination | Synergy score | Most synergistic area score | Method |
| --- | --- | --- | --- |
| Ohmline – Ampicilin (*C. braaki*) | 1.28 | 3.38 | ZIP |
| Ohmline – Ceftriaxone (*C. braaki*) | 7.00 | 10.51 | ZIP |
| Ohmline – Ampicilin (*E. faecalis*) | 9.48 | 26.06 | ZIP |
| Ohmline – Ceftriaxone (C. braaki) | 16.61 | 34.46 | ZIP |
| Ohmline – Ampicilin (*S. aureus*) | 6.98 | 16.04 | ZIP |
| Ohmline – Ceftriaxone (*S. aureus*) | -0,02 | 10.23 | ZIP |

**Supplementary table 3: Viability of microfluidic formulations tested at 24 h.** The same dataset analyzed in figures 3-6 are re-analyzed to calculate the 95% Confidence Interval (CI). 2-4 measurements with 3-9 technical replicas per sample is represented. Outliers are excluded from the analysis.

| *Enterococcus faecalis* | | | | |
| --- | --- | --- | --- | --- |
| **Antibiotic** | | **Ohmline:DMPC (75:25) 0 µg/ml** | **Ohmline:DMPC (75:25) 250 µg/ml** | **Ohmline:DMPC (75:25) 500 µg/ml** |
| **Ampicilin** | **0** | 100 [97,42;102,58] | 90,24 [83,04;97,43] | 60,09 [54,84;65,34] |
|  | **0.33 x MIC** | 73,12 [68,8;77,45] | 38,39 [25,75;51,03] | 40,53 [30,04;51,01] |
|  | **0.41 x MIC** | 69,43 [58,55;80,31] | 16,74 [7,34;26,14] | 5,52 [0,25;10,79] |
|  | **0.5 x MIC** | 19,39 [11,69;27,09] | 5,84 [2,85;8,83] | 1,21 [0;2,46] |
| **Ceftriaxone** | **0** | 100 [97,42;102,58] | 79,97 [76,35;83,59] | 51,02 [44,29;57,75] |
|  | **0.12 x MIC** | 90,16 [88,74;91,59] | 64,24 [61,37;67,1] | 24,09 [18,01;30,17] |
| *Citrobacter braakii* | | | | |
| **Ceftriaxone** | **0** | 100 [99,61;100,39] | 89,79 [85,42;94,17] | 83,69 [81,07;86,31] |
|  | **0.009 x MIC** | 83,77 [81,88;85,66] | 58,87 [55,67;62,06] | 53,33 [53,04;53,62] |
|  | **0.01 x MIC** | 85,75 [82,21;89,29] | 61,54 [56,92;66,16] | 49,59 [46,37;52,82] |
|  | **0.015 x MIC** | 72,83 [69,91;75,74] | 47,44 [43,87;51,00] | 40,66 [37,41;43,91] |
| *Escherichia coli* | | | | |
| **Ampicilin** | **0** | 100 [99,09;100,91] | 83,29 [80,71;85,86] | 82,62 [78,12;87,13] |
|  | **0.86 x MIC** | 73,09 [66,40;79,79] | 60,58 [55,29;65,88] | 60,64 [55,07;66,21] |
| *Klebsiella pneumoniae Kpn10* | | | | |
| **Ciprofloxacin** | **0** | 100 [99,45;100,55] | 89,58 [86,55;92,61] | 78,5 [71,15;85,85] |
|  | **0.125 x MIC** | 71,5 [68,7;74,3] | 63,14 [57,84;68,43] | 50,57 [43,34;57,8] |
| *Klebsiella pneumoniae* clinical sample | | | | |
| **Ciprofloxacin** | **0** | 100 [99,27;100,73] | 86,81 [83,41;90,21] | 71,9 [67,79;76] |
|  | **0.062 x MIC** | 90,01 [88,89;91,13] | 76,44 [70,32;82,55] | 68,43 [66,37;70,49] |
|  | **0.125 µg/ml** | 65,66 [52,46;78,86] | 45,14 [38,3;51,99] | 22,42 [15,64;29,19] |
| *Staphylococcus aureus* | | | | |
| **Ceftriaxone** | **0** | 100 [99,59;100,41] | 90,78 [88,1;93,46] | 85,18 [83,35;87,01] |
|  | **0.0125 x MIC** | 87,16 [84,53;89,8] | 68,62 [63,11;74,13] | 42,37 [33,5;51,24] |
|  | **0.015 x MIC** | 87,60 [84,98;90,21] | 46,52 [40,89;52,15] | 37,35 [32,83;41,87] |
| **Ciprofloxacin** | **0** | 100 [99,49;100,51] | 93,00 [90,26;95,73] | 63,1 [60,18;66,02] |
|  | **0.125 x MIC** | 81,75 [75,84;87,66] | 42,34 [35,3;49,37] | 18,15 [13,98;22,32] |

**Supplementary Figures**

**Supplementary Figure 1. Inhibition of *S. aureus* incubated with DMPC at 500 µg/ml.** The data represent an average of three independent measurements; the error bars indicate the standard deviation of the averaged values. Error bars indicate - SEM. Statistical analysis was performed using t-student test. Statistical significance is indicated as p > 0.05 (ns)

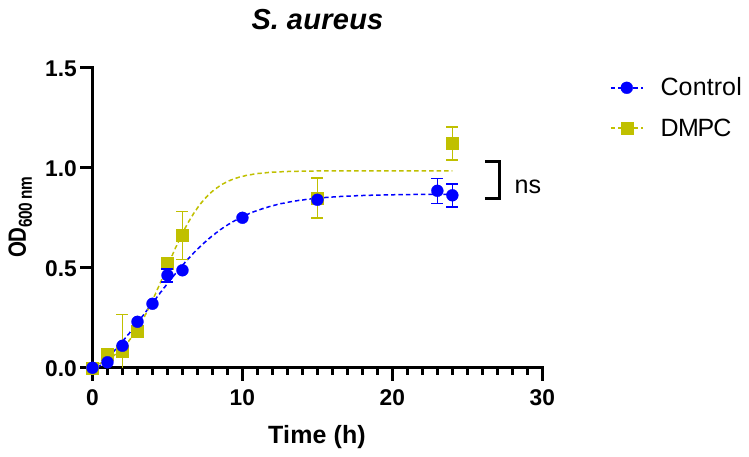

**Supplementary Figure 2. Ohmline biofilm inhibition.** Percentage of biofilm on *S. aureus* in the presence and absence of 500µg/ml of Ohmline. The data represent an average of three independent measurements; the error bars indicate the standard deviation of the averaged values. Error bars indicate - SEM. Statistical analysis was performed using t-student test. Statistical significance is indicated as p < 0.05 (*).

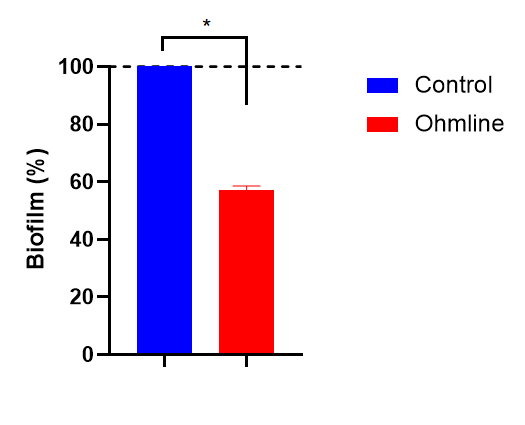

**Supplementary Figure 3. Synergy scores of Ohmline combined with Ampicilin.** 2D charts for Enterococcus *faecalis* (A) y *Sthapylococcus aureus* (B). Obtain with Synergyfinder **https://synergyfinder.aittokallio.group/2025110612260025589/).**

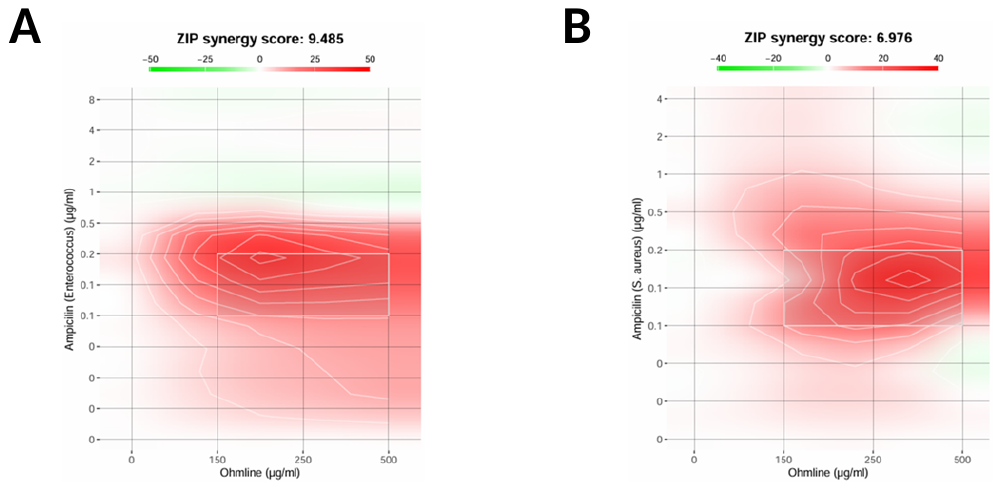

**Fig. S4. Percentage of inhibition of binary combination of Ohmline and commercial antibiotics.**  The data represent an average of three independent measurements. Error bars indicate - SEM. Statistical analysis was performed using one-way ANOVA followed by Tukey’s multiple comparisons test. Statistical significance is indicated as p < 0.05 (*), p < 0.01 (**), and p < 0.0001 (****). The same control growth curve was used for all treatment combinations.

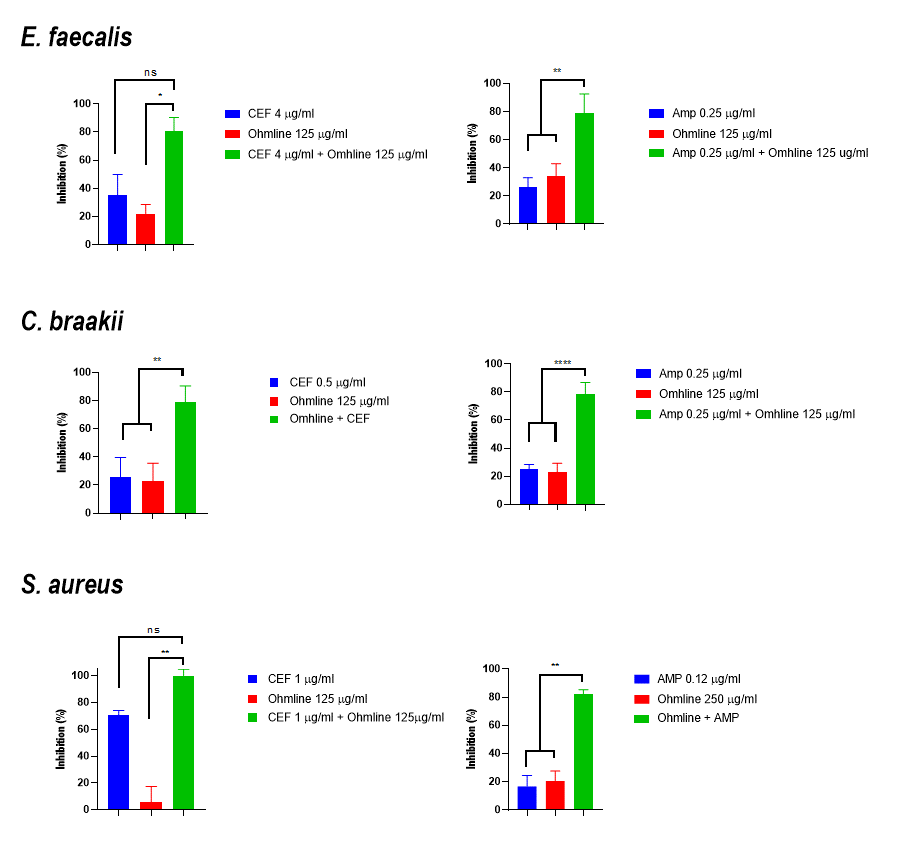

**Supplementary Figure 5. Binary formulations of ampicillin with Ohmline in *Escherichia Coli****.* Optical density was represented as a function of antibiotic concentration and a non-linear regression fit (variable slope) was calculated. The data represent an average of three independent measurements; the error bars indicate the standard deviation of the averaged values.

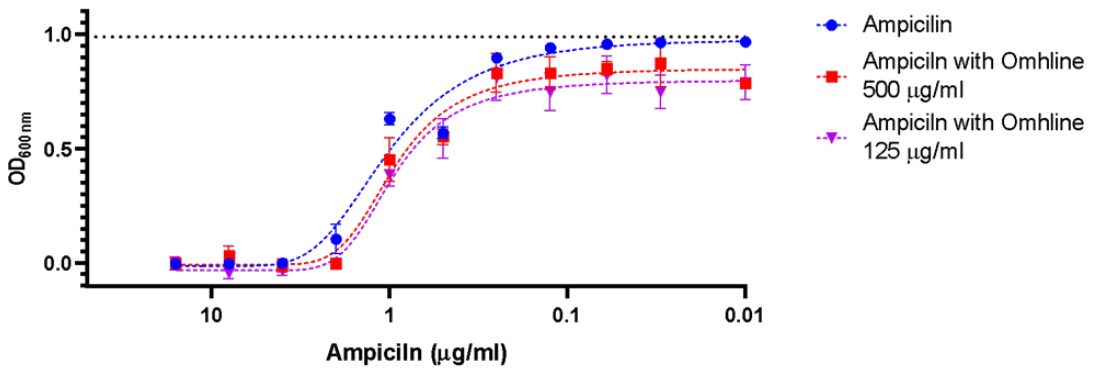

**Fig. S6. DLS measurement.** Diameter size distribution by number is represented. A) Nanoparticles of empty Ohmline:DMPC (75:25); B) Nanoparticles of Ohmline:DMPC (75:25) with Ampicilin; C) Nanoparticles of Ohmline:DMPC (75:25) with Ciprofloxacin; D) Nanoparticles of Ohmline:DMPC (75:25) with Ceftriaxone. Three measurements of the same sample are represented

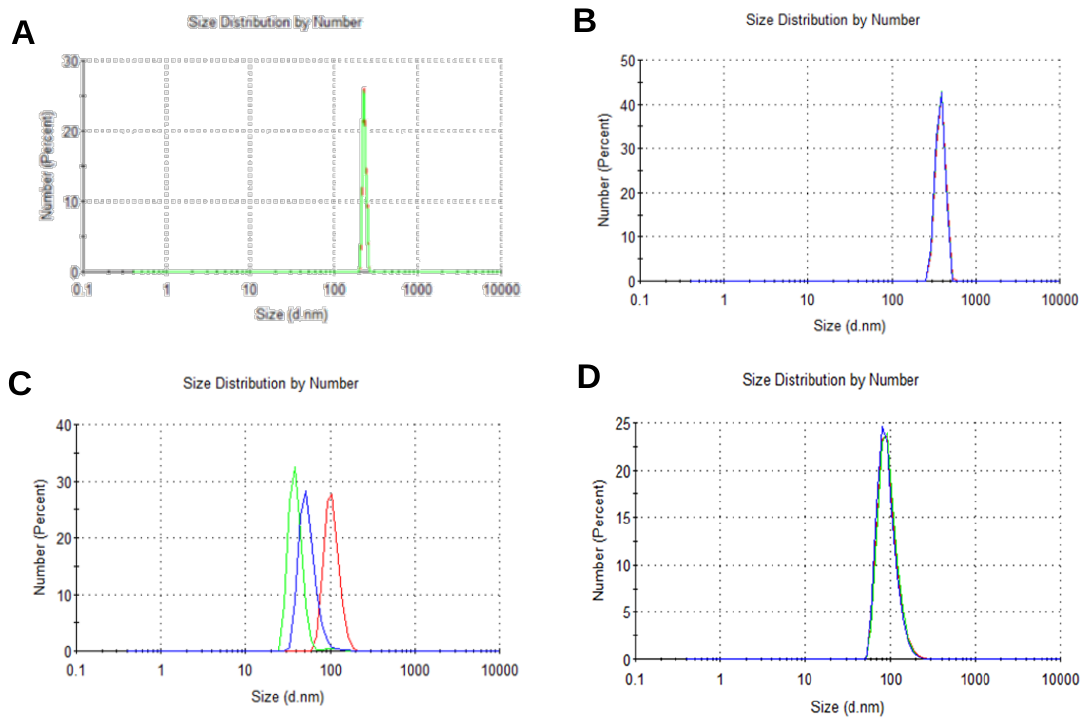

**Supplementry Figure 7. Z potential distribution of lipid nanoparticles**. A) Nanoparticles of empty Ohmline:DMPC (75:25); B) Nanoparticles of Ohmline:DMPC (75:25) with ciprofloxacin; C) Nanoparticles of Ohmline:DMPC (75:25) with ceftriaxone. Three measurements of the same sample are represented.

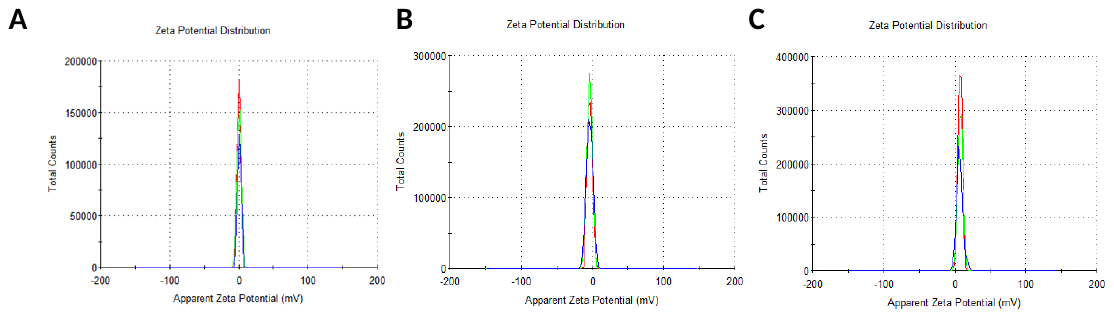

**Fig. S8. Effect of encapsulation of antibiotics on Ohmline nanoparticles at 250 µg/m concentration on the growth of *Enterococcus faecalis*.**  The growth curves obtained with ampicillin (**A-C**) or ceftriaxone **(D)** encapsulated on Ohmline:DMPC (75:25) 250 µg/ml. The data represent an average of three independent measurements with 3-4 technical replicas with ampicillin and 3-7 replicas with ceftriaxone; the error bars indicate the negative SEM. An ordinary one-way ANOVA with Tukey’s multiple comparisons test was performed. Significant differences are represented as: * = pvalue < 0.05; ** pvalue < 0.01; *** = pvalue <0.001; **** = pvalue <0.0001. For each antibiotic measurement, the control curve is used in all concentrations, and is the same as the data in Fig 3.

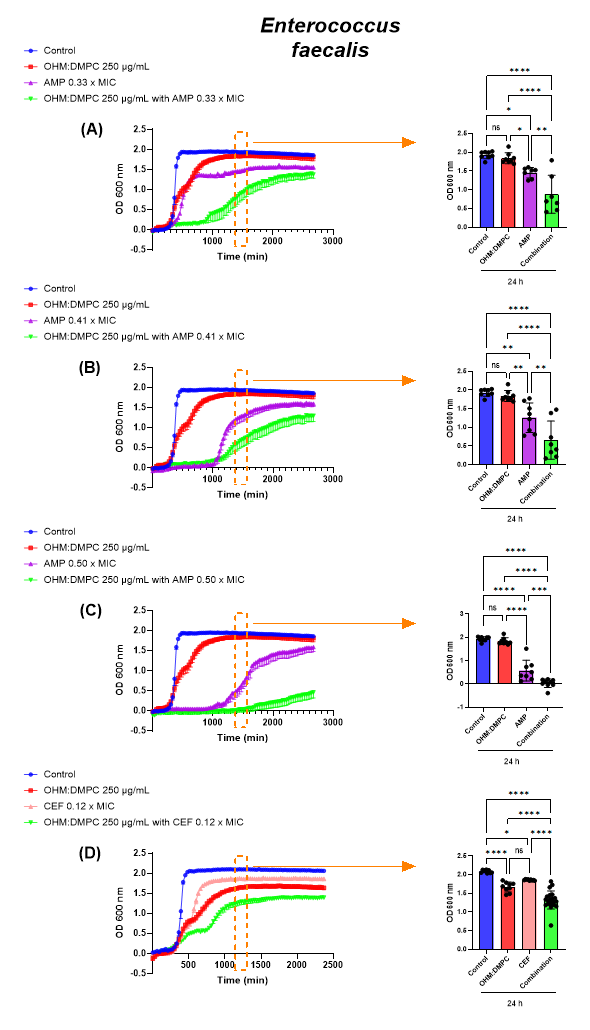

**Supplementary Figure 9**. Effect on the growth of *Escherichia coli* in the presence of ampicillin encapsulated in Ohmline:DMPC (75:25). The data represent an average of three independent measurements; the error bars indicate the standard deviation of the averaged values. The same control curve is used for all concentrations.

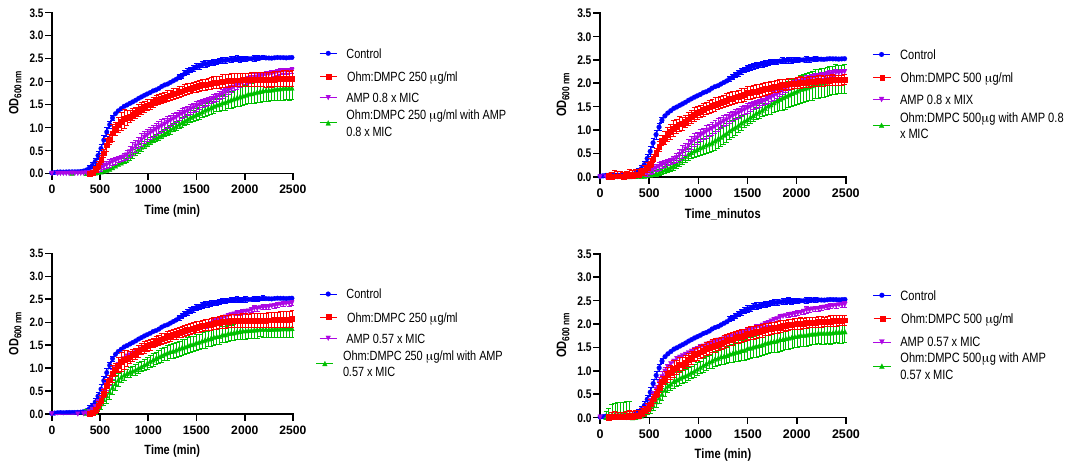
